# Sustained Lysosomal Delivery of Enhanced Cy3-Labeled Acid Nanoparticles Restores Lysosomal pH in Retinal Pigment Epithelial Cells and Astrocytes

**DOI:** 10.1101/2025.06.16.659991

**Authors:** Jiaqi Li, Tianchen Wang, Wennan Lu, Davit Jishkariani, Andrew Tsourkas, Simon Kaja, Rohini M. Nair, Joshua L. Dunaief, Claire H. Mitchell

**Affiliations:** Department of Chemistry, School of Arts and Science; Department of Basic and Translational Science, School of Dental Medicine; Chemical and Nanoparticle Synthesis Core, School of Engineering, University of Pennsylvania, Philadelphia, Pennsylvania, USA United States; Department of Bioengineering, School of Engineering, University of Pennsylvania, Philadelphia, Pennsylvania, USA United States; Department of Ophthalmology, Perelman School of Medicine, University of Pennsylvania, Philadelphia, Pennsylvania, USA United States; Department of Physiology, Perelman School of Medicine, University of Pennsylvania, Philadelphia, Pennsylvania, USA United States; Ophthalmology and Molecular Pharmacology and Neuroscience, Loyola University Chicago, Health Sciences Campus, Maywood, IL, USA

**Keywords:** Lysosomal pH, nanoparticle labeling, nanoparticle trafficking, age dependent neurodegenerations, autophagy, cathepsin D

## Abstract

Lysosomal pH is frequently elevated in age-dependent neurodegenerations like Age-related Macular Degeneration (AMD), Alzheimer’s Disease (AD), and Parkinson’s Disease (PD). Tools that restore lysosomal pH to an optimal acidic range could enhance enzymatic degradation and reduce waste accumulation. Acidic nanoparticles offer a promising strategy for restoring lysosomal function, but accurate tracking of organelle delivery and long-term retention is needed to optimize dosage. To improve detection and enhance delivery, nanoparticles were synthesized from Poly(D,L-lactide-co-glycolide) (PLGA) polymers covalently linked to the fluorescent Cyanine3 amine (Cy3) probe. Nanoparticle concentration and loading times were optimized to achieve >90% delivery to lysosomes in cultured induced pluripotent stem cell-derived retinal pigment epithelial (iPS-RPE) cells. Uptake was heterogeneous, varying between adjacent cells. Once loaded into lysosomes, the nanoparticles were stably retained, with no detectable changes in concentration, distribution, or size for at least 28 days. iPS-RPE cells internalized more nanoparticles than the ARPE-19 cell line or mouse optic nerve head astrocyte cultures. Functionally, PLGA nanoparticles restored an acidic pH and cathepsin D levels in compromised lysosomes. In summary, Cy3-PLGA nanoparticles enabled improved tracking and long-term delivery to lysosomes, supporting future *in vivo* applications to restore lysosomal pH in aging and degenerating tissues.

**Graphical Abstract:** Increased lysosomal pH reduces degradative enzyme efficiency and contribute to age-dependent neurodegeneration. This study describes synthesis of nanoparticles to restore an acidic lumen and degradative function. Nanoparticles were optimized for lysosomal delivery to astrocytes and iPS-derived retinal pigmented epithelial (RPE) cells. The fluorescent marker Cy3 was covalently bound to polymers for improved tracking to lysosomes. Particles were stably retained inside the lysosomal lumen for at least 28 days. Nanoparticles restored pH to compromised lysosomes to baseline levels and increased active Cathepsin D. The improved design will aid *in vivo* tracking and repair in models where lysosomal alkalinization contributes to pathology. Created in BioRender. Mitchell, C. (2025) https://BioRender.com/8hvj96m.

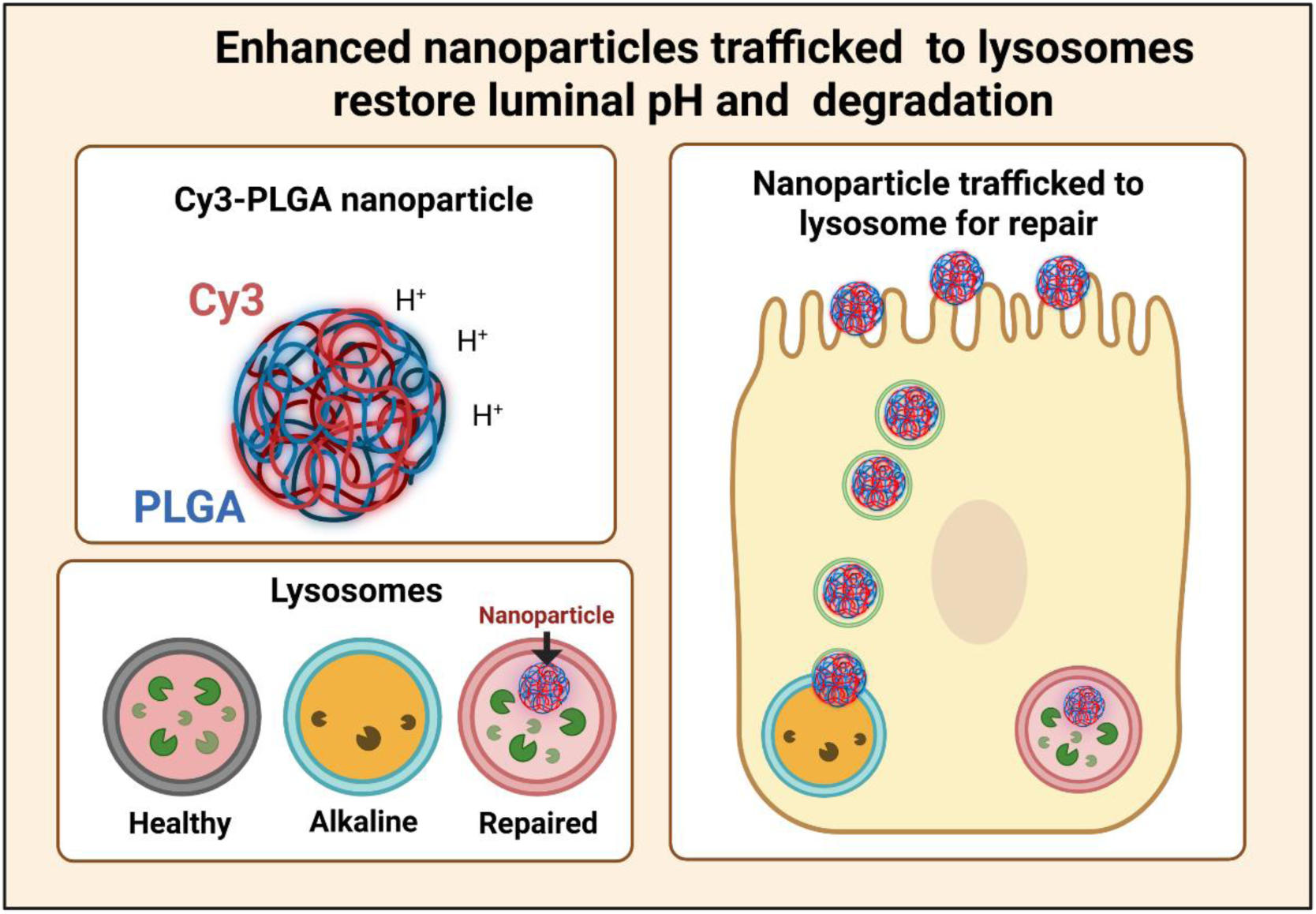

**New and Noteworthy:** Tools that restore acidic pH in compromised lysosomes can enhance autophagy and waste degradation in degenerative disorders marked by excessive accumulation. Here, we describe the novel synthesis of lysosome-targeted nanoparticles composed of PLGA polymers covalently bound to Cy3 fluorescent dye. These Cy3-PLGA nanoparticles enabled improved tracking of lysosomal delivery and demonstrated sustained long-term retention within lysosomes, supporting their potential for future applications to restore lysosomal pH in aging and degenerating tissues.

## Introduction

The decline in degradative function of lysosomal enzymes is a key contributor to waste accumulation and pathology in age-related degenerations [1, 2]. Many lysosomal enzymes are pH dependent, with optimal activity occurring when luminal pH levels are below 5.5 [3, 4]. The pH of the lysosomal lumen is regulated by a complex network of membrane transporters, whose activity can be modulated by second messengers and whose expression is controlled by transcription factors such as TFEB [5, 6]. While this tight feedback system usually meets the cell’s degradative needs are met, even modest elevations in luminal pH found with aging and disease can significantly impair enzyme activity. Therefore, tools that restore an acidic pH to compromised lysosomes may help reduce the accumulation of waste found in many age-dependent neurodegenerative diseases [7–9].

Pharmacological strategies aimed at restoring an acidic lysosomal pH and improving degradative function have shown promise, including the repurposing of FDA-approved drugs, using strategies originally based on pathways that enhanced V-ATPase activity in the plasma membrane [10–13]. While these drug-based approaches have considerable potential, nanoparticles delivered directly to the lysosome could avoid non-specific side effects. Ideally, these nanoparticles would slowly release protons over time, providing sustained acidification. In addition to their therapeutic applications, acidic nanoparticles can serve as valuable tools to directly test whether lysosomal alkalinization drives pathology. To maximize their potential, accurate tracking of lysosomal delivery and long-term retention is essential.

Nanoparticles composed of biodegradable non-toxic polymers such as poly(D,L-lactide) (PLA) and poly(D,L-lactide-co-glycolide) (PLGA), are widely used for controlled-release drug delivery [14, 15]. Degradation of the polymers generates acidic monomers; while the resulting acidification is usually considered detrimental [16], we hypothesized that it could restore an acidic pH to compromised lysosomes if the nanoparticles were delivered to and retained within the lysosomal lumen [17]. PLA polymers, composed of lactic acid, and PLGA polymers, composed of lactic and glycolic acid, have pKa values around 3.8 [18], making them well-suited to buffer lysosomal pH to the normal level of 4.5-4.8. Although rapid escape of PLGA nanoparticles from endo-lysosomal compartments into the cytosol has been reported [19], modulation of nanoparticle size and surface charge is predicted to enhance retention within the lysosomal lumen, but this needs to be evaluated.

The use of nanoparticles to delivery therapeutic treatment to the retina has been well documented [20], and retinal pigmented epithelial (RPE) cells provide an excellent model to evaluate the delivery, retention, and efficacy of acidic nanoparticles. RPE cells are among the most phagocytically active cells in the body, engulfing the tips of photoreceptor outer segments daily [21]. Their lysosomes process copious levels of oxidized lipids and retinoids from this phagocytosed material, in addition to autophagic cargo [22]. As RPE cells are post-mitotic, undegraded material accumulates over time. Excessive lipid waste accumulation is linked to several retinal degenerations, and impaired autophagy is increasingly implicated in disease [23, 24], with changes often detected after additional cell stress such as Ca^2+^ or iron overload [25–27]. Of particular relevance is N-retinylidene-N-retinylethanolamine (A2E), a retinoid byproduct known to elevate lysosomal pH in RPE cells [28–31]. Given the association of A2E with Stargardt’s Disease, restoring an acidic lysosomal pH to RPE cells holds therapeutic potential.

Previous studies demonstrated that PLA-based nanoparticles could lower lysosomal pH in the ARPE-19 cell line [17]. While this provided an important proof of concept, the use of ARPE-19 cells has limitations in terms of physiological relevance and long-term tracking. In addition, the use of Nile Red as a fluorescent nanoparticle marker resulted in cytoplasmic diffusion with some later formulations. To address these issues, the current study used nanoparticles synthesized from PLGA polymer covalently bound to the Cyanine3 amine (Cy3) fluorescent marker for improved specificity and sustained tracking. We also evaluated nanoparticle delivery to the lysosomes of iPS-derived RPE (iPS-RPE) cells, which more closely resemble native RPE cells and are preferable for modeling *in vivo* conditions [32–34]. The extended stability of iPS-RPE cells makes them particularly suitable for long-term retention studies. Finally, this study also examined nanoparticle delivery and efficacy to optic nerve head astrocytes; as lysosomal dysfunction in astrocytes is implicated in both retinal and brain degenerations, restoring lysosomal acidity in these cells may offer additional neuroprotective benefits.

## Methods

### Nanoparticle synthesis

Nanoparticle synthesis was performed by the Chemical and Nanoparticle Synthesis Core at the University of Pennsylvania. Synthesis followed established protocols with several modifications [17]. Polymers 203 H and 502 H (Resomer® R 203 H, Poly(D,L-lactide), Resomer® RG 502 H (Poly(D,L-lactide-*co*-glycolide); all Sigma-Aldrich, St. Louis, MO) were each dissolved in 2 mL of dichloromethane (200 mg per polymer). The solution was added to a 2% w/v aqueous solution of polyvinyl alcohol (PVA) and emulsified *via* probe sonication for 1 min in an ice water bath at 10 W. The resulting primary emulsion (12 mL) was transferred into 50 mL of 2% w/v PVA solution and sonicated for 20 seconds at 3 W to form a secondary emulsion. This mixture was stirred overnight to evaporate the organic solvent. Nanoparticles were collected via gradient centrifugation at 5000, 10000, and 15000 RPM for 5 minutes and 24000 RPM for 20 minutes. Pellets were washed with water, freeze-dried overnight with a lyophilizer, and characterized using a Malvern Zetasizer (Malvern Panalytical, Westborough MA). Nanoparticle sizes ranged from 223-277 nm in diameter, calculated assuming spherical geometry using the manufacturer’s software. The yield was approximately 86%. For early experiments, 0.5 mg/ml Nile Red was included in the primary emulsion.

### Binding polymers with Cy3

Cyanine3 amine (Cy3; Lumiprobe, Hunt Valley, MD) was used to fluorescently label nanoparticles. Cy3 was covalently attached to polymers prior to nanoparticle synthesis. A mixture of 5 mg Cy3, 3 g of 502 H polymer, 12.7 mg EDC (Oakwood Chemical, Estill, SC), 7.6 mg NHS, and 30 µL N,N-Diisopropylethylamine in 50 mL methylene chloride was reacted at room temperature for 48 hours. After the reaction, the resulting solution was taken up in a separatory funnel and were washed three times each with water and 1:1 water/methanol. Organic layers were dried over anhydrous sodium sulfate, filtered, and the filtrates concentrated under reduced pressure using a rotary evaporator. The resulting residue was dried under vacuum overnight and used in nanoparticle synthesis. Due to the high fluorescence intensity of these obtained polymers, a 1:4 ratio of Cy3-labeled to unlabeled polymers was used to produce nanoparticles.

### RPE Cell Culture

The human iPS-RPE cells used in this study were generated and well characterized at the Center for Advanced Retinal & Ocular Therapeutics (CAROT), University of Pennsylvania, Perelman School of Medicine, based on a previously published RPE differentiation protocol [26]. Induced pluripotent stem cells (iPSCs) obtained from the Penn iPSC Core (University of Pennsylvania) were cultured in StemMACS iPS-Brew XF medium (Miltenyi Biotec) on Matrigel-coated plates. Cells were maintained at 37°C in a controlled environment of 5% O₂ and 5% CO₂. Differentiation was initiated once flow cytometry confirmed that over 90% of cells expressed pluripotency markers (SSEA-4 and TRA-1-60). The differentiation process utilized a base medium consisting of DMEM/F12 supplemented with 2% B27, 1% N2, penicillin-streptomycin, and GlutaMAX. Cells were incubated at 37°C with 5% CO₂ for 14 days with the following stage-specific supplements: For days 0-1, noggin (50 ng/mL), DKK1 (10 ng/mL), IGF1 (10 ng/mL), nicotinamide (10 mM) were added, with bFGF (5 ng/ml) added after day 3. For days 4-5, Activin A (100 ng/mL), IGF1 and DKK1 were added, while Activin A, SU5402 (5 μM) and VIP (1 nM) were added for days 6-14. From day 15 onward, cells were maintained in RPE medium (DMEM/Ham’s F12 [70:30], 2% B27, GlutaMAX, and penicillin-streptomycin) until RPE cells with characteristic cobblestone morphology appeared. After 35 days, the cells were dissociated using Accutase/DNase 1 (Sigma-Aldrich) for 30 minutes and subcultured onto Matrigel-coated plates. The subculture medium consisted of TheraPEAK X-VIVO 10 serum-free hematopoietic cell medium supplemented with 100 µg/mL Antibiotic-Antimycotic (Gibco/ThermoFisher, Waltham, MA) and the ROCK inhibitor thiazovivin (5 μM; Tocris Biosciences). Thiazovivin was removed after overnight attachment, and the medium was refreshed every 2-3 days thereafter. The iPSC-derived RPE cells were characterized for mature RPE-specific markers following one month in culture before being used for further experiments. Genotyping revealed heterozygosity for AMD-associated risk alleles in ARMS2 (rs10490924: G/T) and CFH (rs1061170: C/T). Each of these alleles confers a 2.5-fold increased risk of developing age-related macular degeneration (AMD). ARPE-19 cells (ATCC, Manassas, VA,) were cultured in a 1:1 Dulbecco’s modified Eagle medium (DMEM 10-017-CV, Corning) and Ham’s F12 medium with 10% fetal bovine serum (FBS, 35-010-CV, Corning) and 100 µg/ml penicillin/streptomycin (Pen/Strep, Gibco).

### Human ONH astrocyte isolation and culture

Human optic nerve head astrocytes (hONHA) were prepared essentially as described previously [35]. Briefly, donor eyes were obtained via Eversight Eye Bank (Chicago, IL) within 24 hours postmortem. Globes were disinfected with Betadine, dissected under a microscope (M165C, Leica Microsystems, Buffalo Grove, IL), and ONH explants cultured in Ham’s F-10 media with 10% heat-inactivated fetal bovine serum (Gemini Bio, West Sacramento, CA), 2 mM L-glutamine and 0.1 mg/mL Pen/Strep (ThermoFisher). A 50% media change was performed every 2 - 3 days for 8 - 10 weeks. To obtain hONHA cultures, cells were resuspended and maintained for 3 days in serum-free astrocyte basal media (Alkali Scientific, Ft. Lauderdale, FL) before supplementing with 10% FBS. Identity of hONHA was confirmed by immunoblotting; hONHA were positive for glial fibrillary acidic protein, laminin, but negative for alpha smooth muscle actin (not shown). The Study was conducted in compliance with the tenets of the Declaration of Helsinki, and reviewed by the Institutional Review Board for the Protection of Human Subjects of Loyola University Chicago (#LU217139).

### Nanoparticle delivery

iPS-RPE cells were plated on Geltrex-coated coverslips in 24-well plates and grown to confluence. Cy3-PLGA nanoparticles (0.5-2.0 mg/ml) in growth media were sonicated for 10 min (B200 Ultrasonic Cleaner, Branson; Input Power: 27W; Output Power: 19W) before application. Cells were incubated at 37°C for the indicated time, then washed with Mg^2+^/Ca^2+^-free Dulbecco’s Phosphate Buffered Saline (DPBS; ThermoFisher), replenished with fresh media and returned to the incubator. Cells were typically examined after a 2-hour “chase”. For imaging, cells were stained with 200 nM LysoTracker Green DND-26 (ThermoFisher) and 1 μM Hoechst 33342 for 15 minutes, washed, and imaged using Live Cell Imaging Solution (ThermoFisher). Imaging was conducted using a Nikon Eclipse microscope with NIS Elements Imaging software (All Nikon USA, Melville NY). Filter sets used were Hoechst 33342: Excitation/Emission (Ex/Em) 385/460 nm, LysoTracker Green/Bodipy Pepstatin A: Ex/Em 485/520 nm, Cy3/LysoTracker Red/Nile Red: Ex/Em 560/602 nm. Confirmation was performed using a Nikon Eclipse Ti2-microscope and Crestor spinning disk confocal system.

### Uptake time course and cell type comparison

Cells were incubated with 1 mg/ml Cy3-PLGA nanoparticles for 24 hrs, washed 3x and returned to growth medium. Imaging occurred on days 1, 7, 21 and 28 days after nanoparticle removal. Data represent the mean of 3 measurements per field, with 6 fields per time point. iPS-RPE, ARPE-19 and mONH astrocytes were grown for 5 days, exposed to nanoparticles and imaged similarly.

### Image analysis

Nanoparticle uptake was quantified manually at x40 from a mean of 24 cells per field using Hoechst staining. Percentage uptake was calculated as 100*(# of cells with red particles/total cells). For particle quantification and colocalization, a x100 objective was used and images analyzed in ImageJ [36]. The “Analyze Particles” function detected particles >10 pixels. Colocalization was defined as the presence of both green and red signals. Colocalization percentages were calculated as 100*(# of particles with red and green)/(# red particles). The % of lysosomes with nanoparticles was defined as 100*(# of particles with red and green)/(# green particles).

### Lysosomal pH measurement

Lysosomal pH was measured using the LysoSensor Yellow/Blue DNS 160 probe (#L7545, ThermoFisher) as detailed [37]. Cells were cultured in 96-well black plates with clear bottoms using appropriate media until confluent (3-4 days). Unlabeled PLGA or PLA nanoparticles were added to media at 2 mg/ml and sonicated for 10 minutes, then added to cells as appropriate, with cells grown for a further 5-7 days. After washing, and incubation with tamoxifen for 30 minutes at 37°C, cells were incubated in isotonic solution with 4 µM LysoSensor Yellow/Blue DND 160 for 8 minutes at 37°C. Isotonic solution contained: NaCl, 105 mM; KCl, 5 mM; HEPES-Acid, 6 mM; Na-HEPES, 4 mM; NaHCO_3_, 5 mM; mannitol, 60 mM; glucose, 5 mM; MgCl_2_, 0.5 mM; CaCl_2_, 1.3 mM; pH, adjusted to 7.4; osmolality, 300 mOsm. Cells were washed 3x in isotonic solution, then imaged in a Fluoroskan 96-well plate reader (ThermoFisher). Lysosomal pH was determined from the ratio of light excited at 340nm versus 380nm (>520nm em), with 10 measurements averaged. As calibration to absolute pH levels gave inconsistent results, values are expressed as the 340nm/380nm excitation ratio and normalized to each day’s control value, as levels varied between experimental sets [38].

### Cathepsin D measurement

BODIPY FL-Pepstatin A (ThermoFisher) was used to assess Cathepsin D levels, as adapted from Guha et al. [10]. hONH astrocytes were grown in 8-well chamber slides until confluent (5-12 days), treated overnight with 1 mg/mL nanoparticles, and stained with 1 µg/ml BODIPY FL-Pepstatin A and 25µM tamoxifen for 30 min at 37°C. Imaging was performed with a Nikon Eclipse microscope (Ex/Em 485/520 nm) and analyzed in ImageJ by thresholding and measuring the area above threshold. Data were normalized to daily control means. Costaining with LysoTracker Red (Ex/Em 560/602 nm) confirmed lysosomal localization of the Pepstatin A signal.

### Statistical analysis

GraphPad Prism v10 was used for statistical analysis (GraphPad Software, San Diego, CA). Normality was assessed via Shapiro-Wilk or Kolmogorov-Smirnov tests. ANOVA or non-parametric tests were applied as appropriate. Multiple comparisons were corrected using Tukey’s, Dunnett’s or Dunn’s tests as appropriate. Time trends (Figure 5) were analyzed with a first-order regression using a least-squared approach. Significance was defined as p < 0.05. Data are presented as mean ± standard deviation (SD).

All compounds were from Sigma-Aldrich Corporation (St. Louis, MO), unless otherwise stated.

## Results

### Synthesis of Cy3-PLGA nanoparticles

Nanoparticles were synthesized from PLGA and PLA polymers to optimize uptake into and retention within lysosomes. As nanoparticle size can influence cellular uptake and intracellular trafficking [39], sonication and centrifugation parameters were adjusted to modify particle diameter. Dynamic light scattering analysis using a Malvern Zetasizer revealed nanoparticle diameters ranging from 222 to 277 nm. To provide a more stable fluorescent labeling for intracellular tracking, Cyanine3 amine (Cy3) was covalently conjugated to PLGA polymers (Fig. 1A), and this Cy3-labeled polymer was used to synthesize the nanoparticles (Fig. 1B).

**Figure 1.**
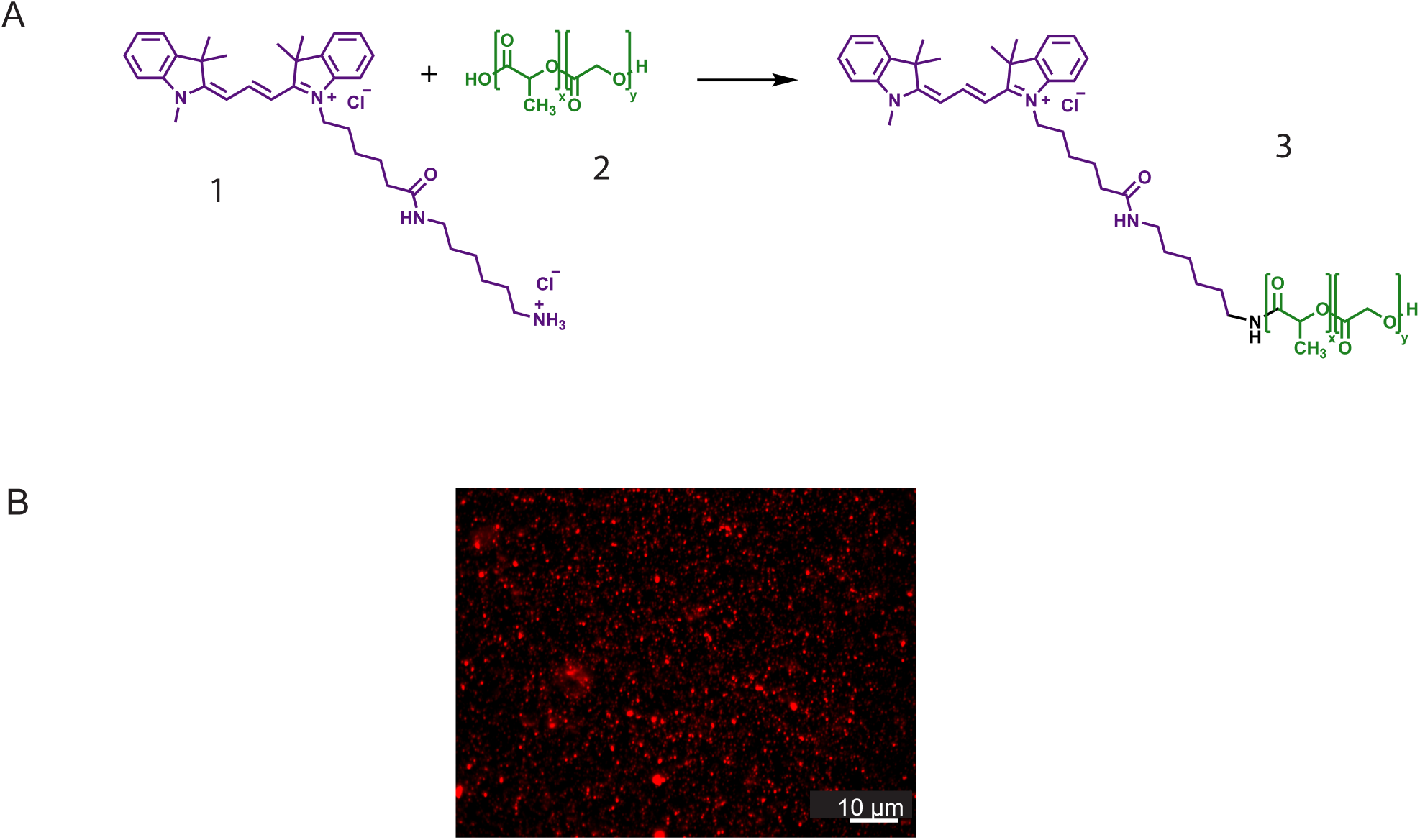
Synthesis of nanoparticles and covalent labeling with Cy3. A. Schema of Cy3 conjugation to PLGA polymers. Poly(D,L-lactide-co-glycolide) (PLGA) polymers are composed of lactic acid and glycolic acid monomers. The amine group of Cyanine3 amine (1) reacts with the carboxylic acid terminus of PLGA (2), forming a stable amide bond and generating Cy3-labeled PLGA polymers (3). B. Visualization of Cy3-labeled PGLA nanoparticles (red).

### Incubation time and delivery of Cy3-PLGA acid nanoparticles to iPS-RPE cells

The bright Cy3 fluorescence of the PLGA-based acid nanoparticles enabled detailed analysis of their lysosomal delivery. Initial experiments investigated the effect of incubation time on lysosomal uptake. Cy3-PLGA nanoparticles were applied at a constant concentration of 1 mg/ml to confluent iPS-RPE monolayers for 1, 2, or 24 hours. Following incubation, cells were washed and an additional two-hour “chase” period was allowed for intracellular trafficking.

After a 1-hour incubation, only a few red fluorescent nanoparticles were detected in iPS-RPE cells (Fig. 2A). A 2-hour incubation led to a moderate increase in Cy3-labeled particles (Fig. 2B), while a 24-hour incubation resulted in a substantially larger rise in the fluorescent signal (Fig. 2C). Quantification confirmed this trend; the mean number of nanoparticles per cell increased from 6 after 1 hour to 17 after 2 hours, and reached 50 after a 24 hour incubation. (Fig. 2D). Likewise, the percentage of cells containing at least one nanoparticle increased with incubation time, from 34% of cells at 1 hour to 73% after 24 hours (Fig. 2E).

**Figure 2.**
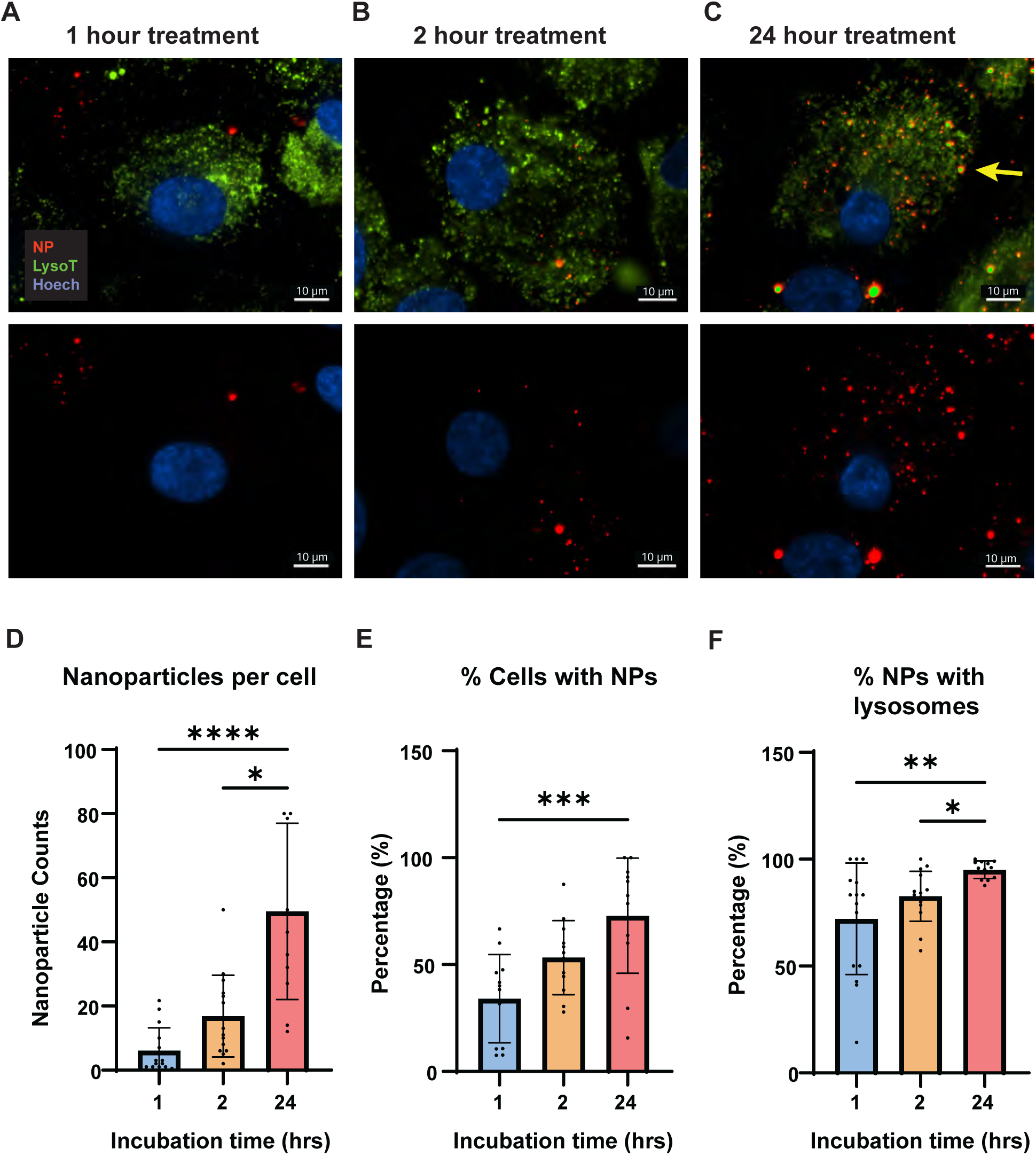
Time-dependent delivery of Cy3-PLGA nanoparticles to lysosomes in iPS-RPE cells. A. Representative images of iPS-RPE cells after a 1-hour incubation with 1 mg/ml Cy3-PLGA nanoparticles (NP, red). Minimal colocalization with LysoTracker Green-labeled lysosomes (LysoT, green) was observed. Nuclei are stained with Hoechst 33342 (Hoech, blue). Top panels show merged images, bottom panels show red and blue channels only. All cells were washed and incubated in a nanoparticle-free medium for 2 hours prior to imaging to provide time for trafficking. B. After a 2-hour incubation, nanoparticle retention increased slightly, and some nanoparticles colocalized with lysosomes (yellow overlap). C. A 24-hour incubation increased nanoparticle uptake and colocalization with lysosomes (yellow, arrow). D. Quantification of nanoparticle count per cell showed a significant increase with longer incubation time. * p=0.0277, **** p<0.0001 (Kruskal-Wallis test with Dunn’s multiple comparisons test; n=12-15 fields from 2 independent experimental trials). Throughout, bars are mean ± SD, dots represent values from individual images. E. The percentage of iPS-RPE cells with at least one nanoparticle also increased with incubation time. *** p=0.0004 (One-way ANOVA with Tukey’s multiple comparisons test, n=12 fields from 2 independent trials). F. The proportion of nanoparticles colocalizing with LysoTracker Green increased with incubation time. * p=0.024, ** p=0.0074 (Kruskal-Wallis test with Dunn’s multiple comparisons test; n=12 fields from 2 independent trials).

To assess the subcellular localization of the Cy3-PLGA nanoparticles, colocalization with LysoTracker Green—a lysosomal marker—was analyzed. Even after 1 hour, more than half of the nanoparticles colocalized with LysoTracker Green (Fig. 2F). As all incubations were followed by a 2-hour rest period without nanoparticles, this indicates that 2 hours was sufficient for the trafficking of nanoparticles to lysosomes. After a 24-hour incubation period, nearly all (95%) of the Cy3-labeled nanoparticles colocalized with lysosomes.

### Nanoparticle concentration and the delivery of Cy3-PLGA acid nanoparticles to iPS-RPE cells

Given the improved detection of the Cy3-PLGA nanoparticles, the effect of nanoparticle concentration on uptake and lysosomal delivery was evaluated. All measurements were performed after a 24-hour incubation followed by a 2-hour chase period, as this produced stable delivery in the preceding experimental set. The number of Cy3-PLGA particles per cell increased with concentration across the tested range of 0.5, 1.0 and 2.0 mg/ml (Fig. 3A-C). At 0.5 mg/ml, a mean of 20 nanoparticles were detected; this increased significantly with 2.0 mg/ml (Fig. 3D). Higher concentrations also increased the percentage of cells containing at least one nanoparticle, with over 80% of cells showing uptake at 2.0 mg/ml (Fig. 3E). Despite this, uptake varied considerably; even at 2.0 mg/ml, adjacent cells displayed markedly different levels of nanoparticle accumulation.

**Figure 3.**
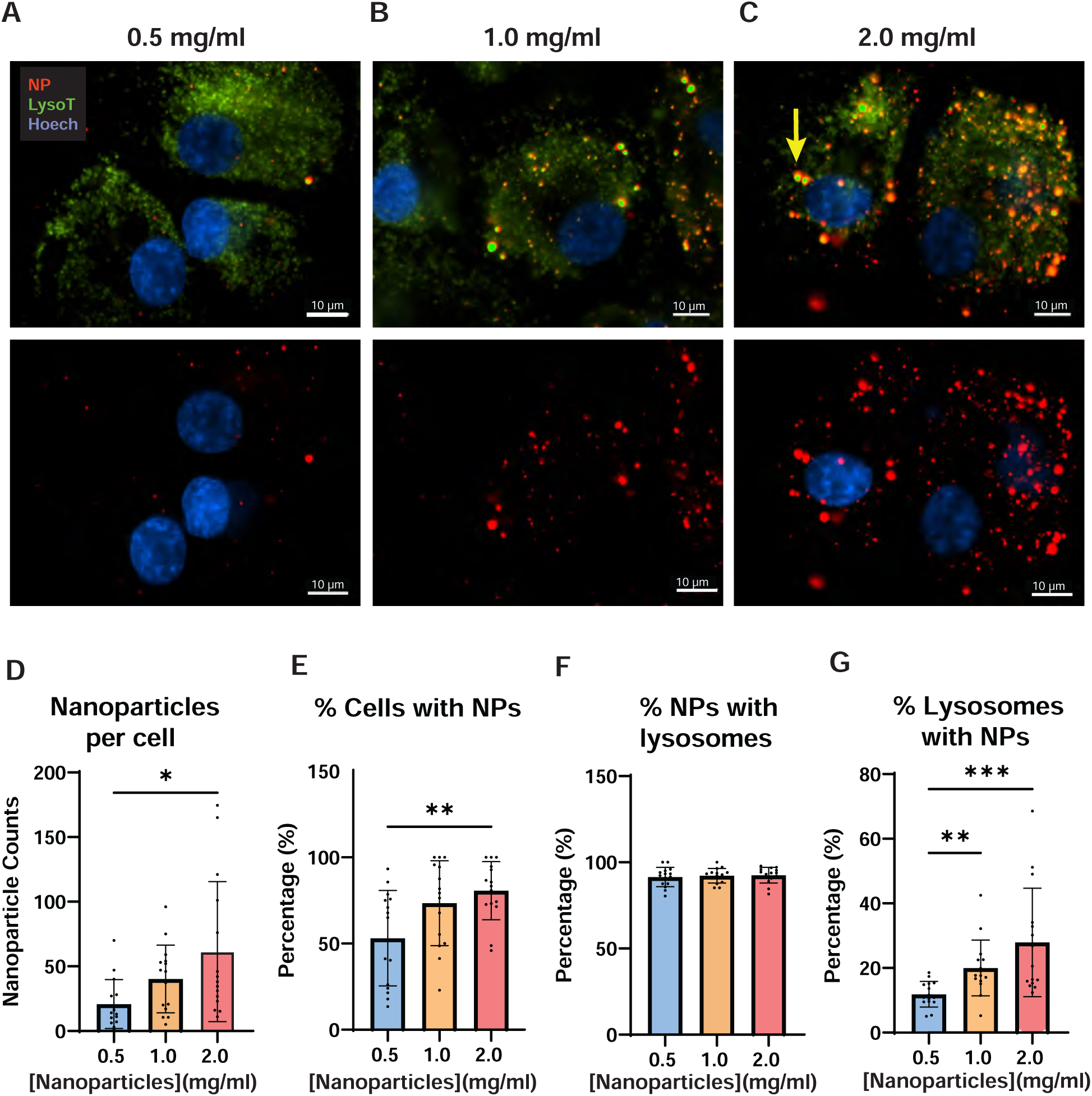
Effect of Cy3-PLGA nanoparticle concentration on accumulation in iPS-RPE cells. A-C. Representative images of iPS-RPE cells following 24-hour incubation with Cy3-PLGA nanoparticles at increasing concentrations (0.5, 1.0 and 2.0 ng/ml). Arrow highlights example of colocalization of LysoTracker (LysoT, green) with Cy3-PLGA nanoparticles (NP, red). Nuclei are stained with Hoechst 33342 (Hoech, blue). Top panels show merged images, bottom panels show red and blue channels only. All cells were washed and incubated in a nanoparticle-free medium for 2 hours prior to imaging D. Nanoparticle count per cell increased significantly with concentration. * p=0.0149 (Kruskal-Wallis test with Dunn’s multiple comparisons test; n=15 fields from 2 trials). E. The percentage of iPS-RPE cells containing at least one nanoparticle also rose with concentration. ** p=0.0069 (One-way ANOVA with Tukey’s multiple comparisons test; n=15). F. The percentage of nanoparticles localized to lysosomes remained constant across concentration (One-way ANOVA with Tukey’s multiple comparisons test; n=15). G. The percentage of lysosomes containing nanoparticles increased significantly with concentration. ** p=0.0043, *** p=0.0005 (Kruskal-Wallis test with Dunn’s multiple comparisons test; n=15).

To determine whether the endocytosed materials typically traffic to lysosomes, nanoparticle localization was assessed via colocalization with LysoTracker Green. At all concentrations, more than 90% of Cy3-labeled nanoparticles colocalized with LysoTracker Green (Fig. 3F). Although LysoTracker is less sensitive to pH changes compared to other probes, the colocalization of green and red fluorescent signals could have reflected acidification of other organelles containing a nanoparticle. However, most LysoTracker-positive organelles did not colocalize with Cy3-PLGA nanoparticles, and the percentage of red-green colocalization increased with loading concentration, from 12% at 0.5 mg/ml to 28% at 2 mg/ml (Fig. 3G).

Confocal imaging confirmed the cell-to-cell variability in nanoparticle uptake, with some cells harboring only a few particles while neighboring cells exhibited substantial accumulation (Fig. S2). While difficult to quantify precisely, this heterogeneity was a consistent and characteristic feature of nanoparticle uptake.

### Sustained retention of Cy3-PLGA nanoparticles in iPS-RPEs

One of the key advantages of iPS-RPE cells is their ability to maintain stable growth for months when cultured in a media that supports differentiation [40]. This makes them preferable to ARPE-19 cells for long-term studies of nanoparticle delivery and stability. To assess long-term retention, iPS-RPE cells were incubated with 1 mg/ml Cy3-PLGA nanoparticles for 24 hours, followed by analysis of nanoparticle characteristics at 1, 7, 14, 21, and 28 days post-loading. Cells retained a healthy appearance throughout the 28-day period, with no noticeable morphological changes observed following nanoparticle exposure (Fig. 4A–C).

**Figure 4.**
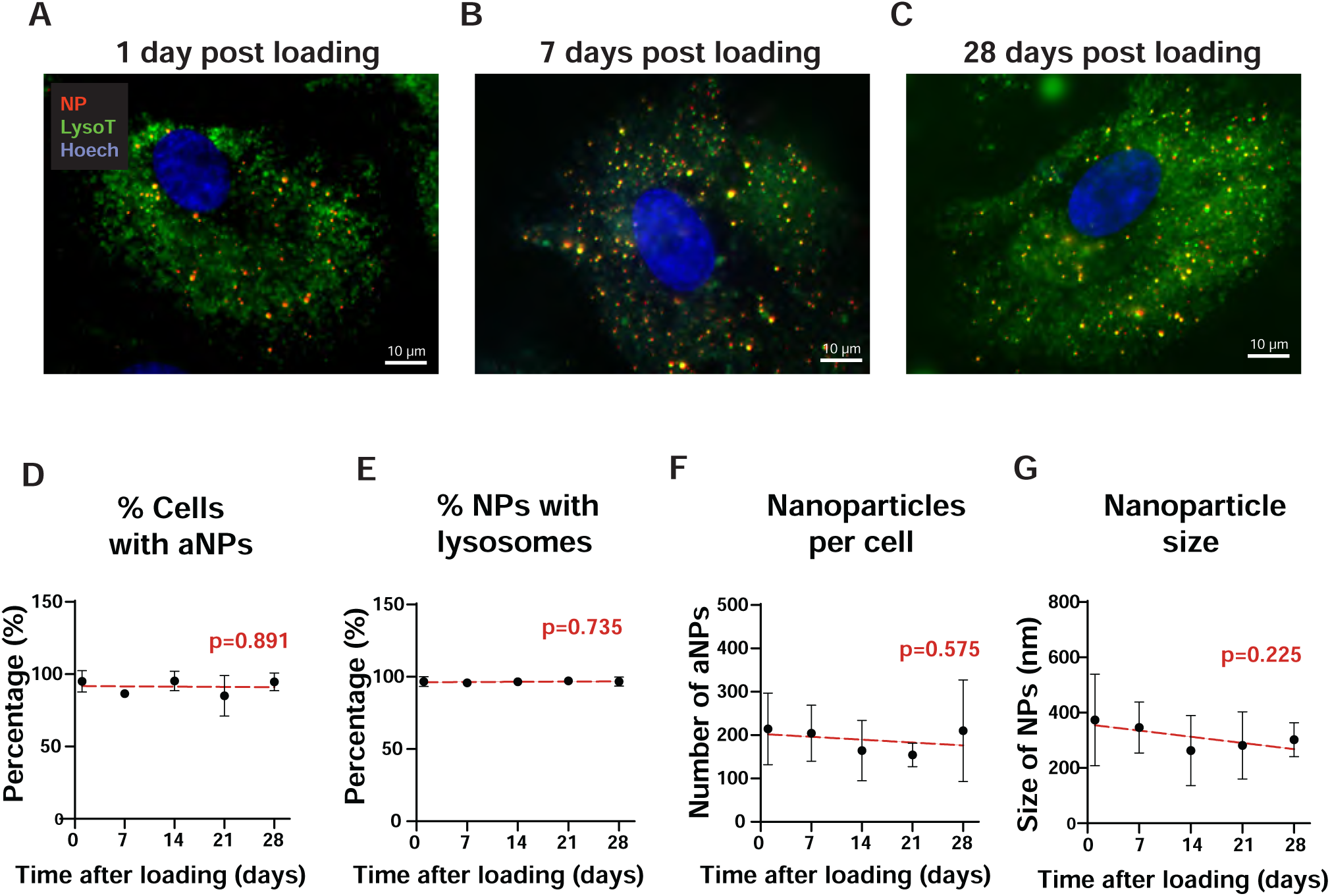
Retention of nanoparticles over time. A-C. Representative images of iPS-RPE cells 1, 7 and 27 days after a 24-hour exposure to 1.0 mg/ml Cy3-PLGA nanoparticles (NP, red); LysoTracker Green (LysoT, green); Hoechst 33342 (Hoech, blue). Yellow staining indicates overlap of Cy3-labeled PLGA nanoparticles with LysoTracker Green. D-G. Quantification of nanoparticle retention over time. D. % of cells containing nanoparticles; E. % of nanoparticles colocalized with lysosomes; F. Nanoparticle count per cell; G. Mean size of nanoparticles per cell. Each data point represents mean ± SD from 6 fields (Days 1-21) or 3 fields (Day 28). Lines represent linear regression fits; red text indicates p-values for slope comparisons with zero slope.

At each time point, three key metrics were evaluated: the percentage of cells containing at least one nanoparticle, the percentage of nanoparticles colocalized with LysoTracker Green, and the mean number of nanoparticles per cell. Additionally, particle size was measured to assess whether nanoparticles shrank over the 28-day period. Data were analyzed using first-order linear regression, and the slopes were tested for statistical significance to determine whether any changes occurred over time.

No significant changes were observed in the percentage of cells containing nanoparticles (Fig. 4D) or in the proportion of nanoparticles colocalized with lysosomes (Fig. 4E) across the 28-day period. Although there was a slight downward trend in the number of nanoparticles per cell, the change was not statistically significant (Fig. 4F). Similarly, a modest decrease in particle size over time was observed, but this trend also failed to reach statistical significance (Fig. 4G). Collectively, these results indicate that the Cy3-PLGA nanoparticles remained stable within iPS-RPE cells for at least 28 days, supporting their potential for long-term restoration of acidic lysosomal pH.

Notably, the average measured diameter of fluorescent particles within cells was 305 ± 37 nm. Although this measurement is limited by the spatial resolution of light microscopy and should be interpreted cautiously, it closely matches the 278 nm mean diameter determined by Zetasizer analysis of the nanoparticle batch used in the study. This similarity supports the accuracy and reliability of intracellular detection.

### Comparison of nanoparticle uptake by different cell types

RPE cells are among the most phagocytically active cells in the body, routinely engulfing the tips of photoreceptor outer segments each day [21, 41]. While ARPE-19 cells were the standard cell model for many years, recent advances in stem cell technology have led to the development of iPS-RPE cell lines, which more closely resemble RPE cells *in vivo*. Astrocytes also exhibit phagocytic activity and take up debris such as amyloid beta to enhance the environment surrounding neurons. Our prior characterization of astrocytes isolated from the mouse optic nerve head supports their use in this study [42].

Confluent cultures of iPS-RPE, ARPE-19, and primary astrocytes were exposed to 1 mg/ml Cy3-PLGA nanoparticles for 24 hours. Following a 2-hour chase period, nanoparticle distribution was assessed. All three cell types internalized nanoparticles (Fig. 5A–C), and quantification revealed no significant difference in the percentage of cells that internalized at least one nanoparticle (Fig. 5D). However, the number of nanoparticles internalized per cell varied across cell types, with iPS-RPE cells showing markedly higher uptake (Fig. 5E).

Distinct differences in nanoparticle uptake patterns and cellular morphology were also observed. iPS-RPE cells exhibited a higher lysosome density, indicated by green puncta in LysoTracker Green-labeled cells. In contrast, ARPE-19 cells showed fewer lysosomes, though some appeared to associate with larger Cy3-positive regions, potentially reflecting either the aggregation of multiple nanoparticles within individual lysosomes or preferential uptake of larger nanoparticles. Astrocytes were larger than either RPE cell type and internalized fewer nanoparticles per cell, resulting in a lower overall density. Additionally, nanoparticles appeared smaller within astrocytes. These observations suggest that nanoparticle uptake is influenced by cell type–specific properties, rather than occurring randomly.

### PLGA-based nanoparticles restore lysosomal pH

Many lysosomal degradative enzymes function optimally at low pH [3]. However, lysosomal pH can rise with age and disease [43], and strategies to restore acidic conditions may hold therapeutic potential. To evaluate whether acid nanoparticles could help normalize lysosomal pH, we used the ratiometric indicator LysoSensor Yellow/Blue DND-26. Although somewhat sensitive to experimental conditions, this dye provides reliable results with appropriate precautions [38]. The overall goal of this project is to restore an acidic pH to alkalinized lysosomes, and previous studies have shown that some interventions are more effective when lysosomal pH is already elevated [37]. To induce a moderate rise in lysosomal pH, cells were treated with tamoxifen, which reliably increases organelle pH via proton transport across membranes, independent of estrogen receptor signaling [44, 45], and is more consistent in this regard than chloroquine or bafilomycin A.

iPS-RPE cells were pretreated for 5 days with PLGA or PLA nanoparticles; as the LysoSensor readout is fluorescent, nanoparticles synthesized in parallel from polymers not bound to Cy3 were used. After a 30-minute tamoxifen treatment, the lysosomal pH was measured. Tamoxifen significantly elevated lysosomal pH, but this increase was attenuated in cells pretreated with PLGA nanoparticles (Fig. 6A). In contrast, pretreatment of cells with PLA nanoparticles did not significantly affect pH levels.

**Figure 5.**
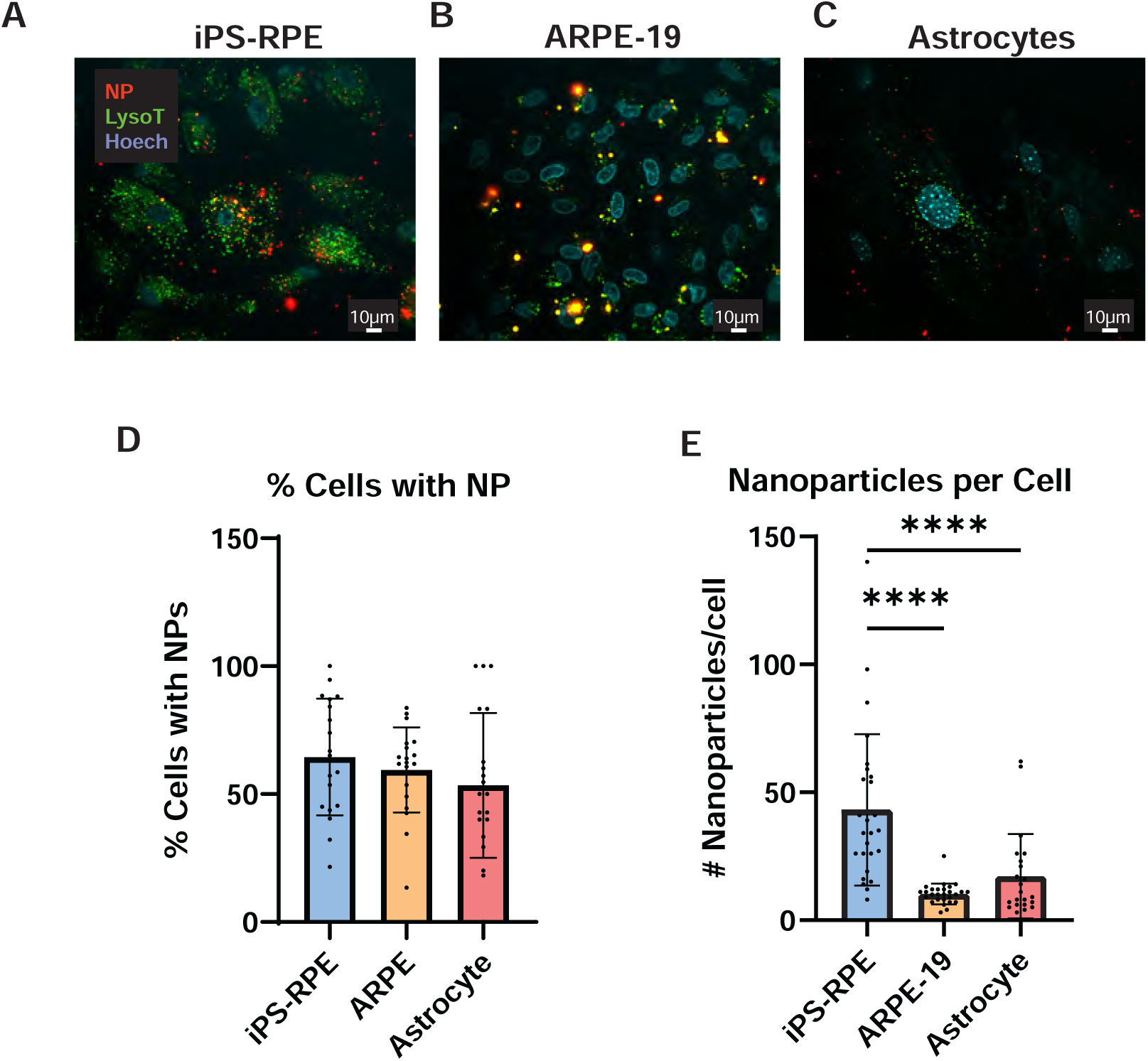
Nanoparticle uptake across different cell types. A-C. Representative images of nanoparticle uptake by iPS-RPE cells, ARPE-19 cells and primary mouse optic nerve head astrocytes (ONHA), respectively. All incubated with 1 mg/ml Cy3-PLGA nanoparticles for 24 hrs, followed by a 2-hour washout. NP: Cy3-labeled PLGA nanoparticle; LysoT: LysoTracker Green; Hoech: Hoechst 33342. D. Quantification shows the percentage of cells with at least one nanoparticle was similar in all three cell types. (One-way ANOVA; n=19-20). E. There was significantly greater nanoparticle uptake in iPS-RPE cells compared to ARPE-19 and astrocytes. **** p<0.0001 (Kruskal-Wallis test with Dunn’s test; n=22-27 fields).

**Figure 6.**
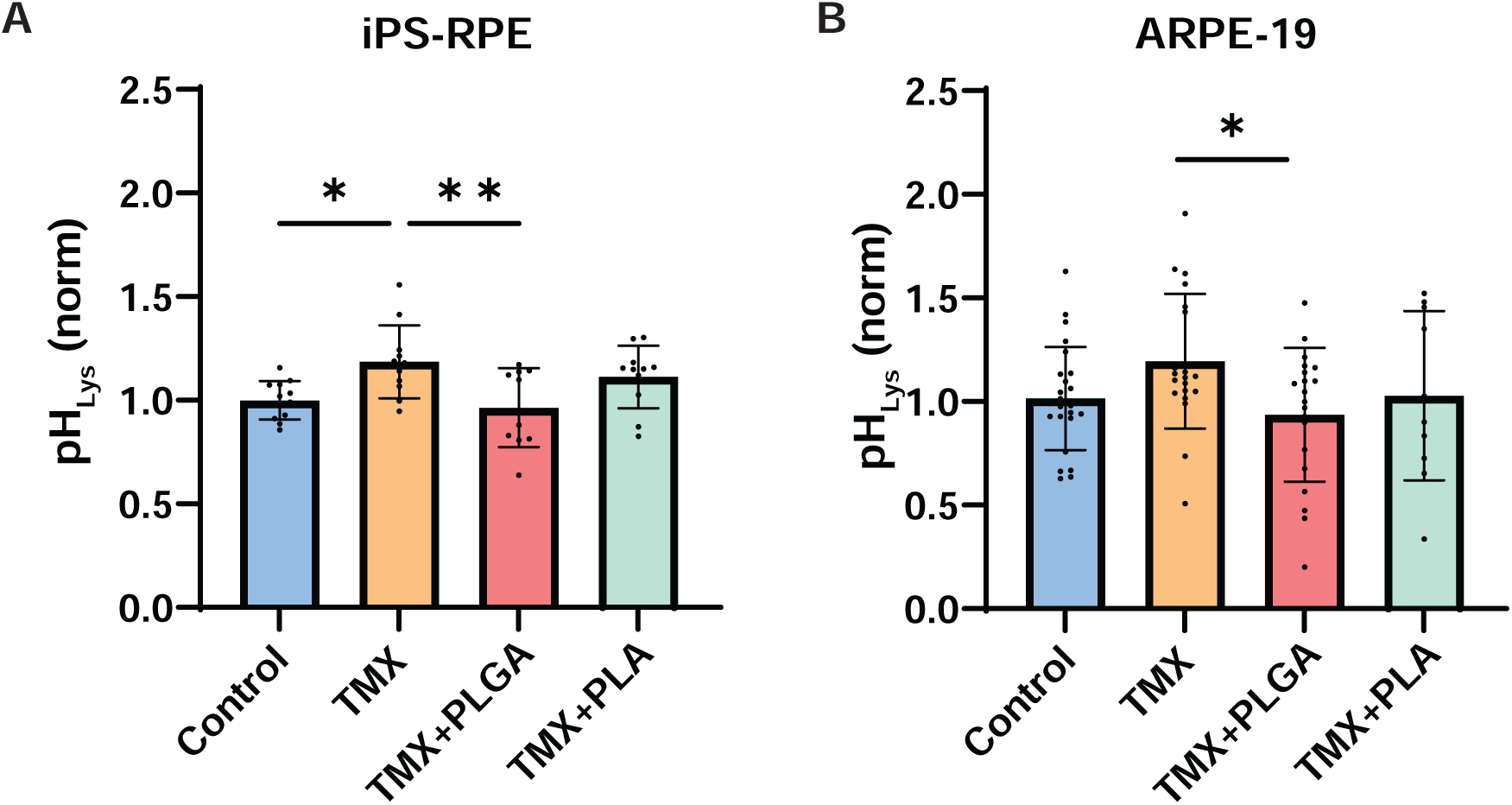
Acidic nanoparticles restore lysosomal pH in compromised iPS-RPE and ARPE-19 cells. A. Lysosomal pH in iPS-RPE cells was returned to baseline levels with PLGA nanoparticle treatment. Cells were pretreated with 2 mg/ml unlabeled PLGA- or PLA-based nanoparticles for 5 days before 30 min tamoxifen exposure (TMX, 10 µM). * p=0.0178, ** p=0.0062 (One-way ANOVA with Dunnett’s test; n=10-12). Values normalized to daily controls based on ratiometric imaging with LysoSensor Yellow/Blue (pH_Lys_ = 340/ 380 nm excitation, >527 nm emission). B. Similar results were observed in ARPE-19 cells, where only PLGA nanoparticles reversed tamoxifen-induced pH elevation. *p=0.0308 (Dunnett’s test; n=10-25 from 2-3 trials).

The ability of PLGA- or PLA-based nanoparticles to restore an acidic pH to lysosomes of ARPE-19 cells was also examined. As observed with iPS-RPE cells, pretreatment with PLGA nanoparticles reduced the lysosomal pH following tamoxifen exposure (Fig. 6B), whereas pretreatment with PLA nanoparticles had no significant effect, mirroring the response in iPS-RPE cells.

### Nanoparticles restore lysosomal activity

The functional effects of acidic nanoparticles on astrocytes were also investigated. Primary cultures of human optic nerve head astrocytes were used due to their favorable growth characteristics and increased physiological relevance [46]. Lysosomal pH measurement confirmed that tamoxifen treatment increased lysosomal pH in the astrocytes while PLGA nanoparticle treatment partially reversed this alkalinization, resulting in a mean decrease of 37% (Fig. 7A).

**Figure 7.**
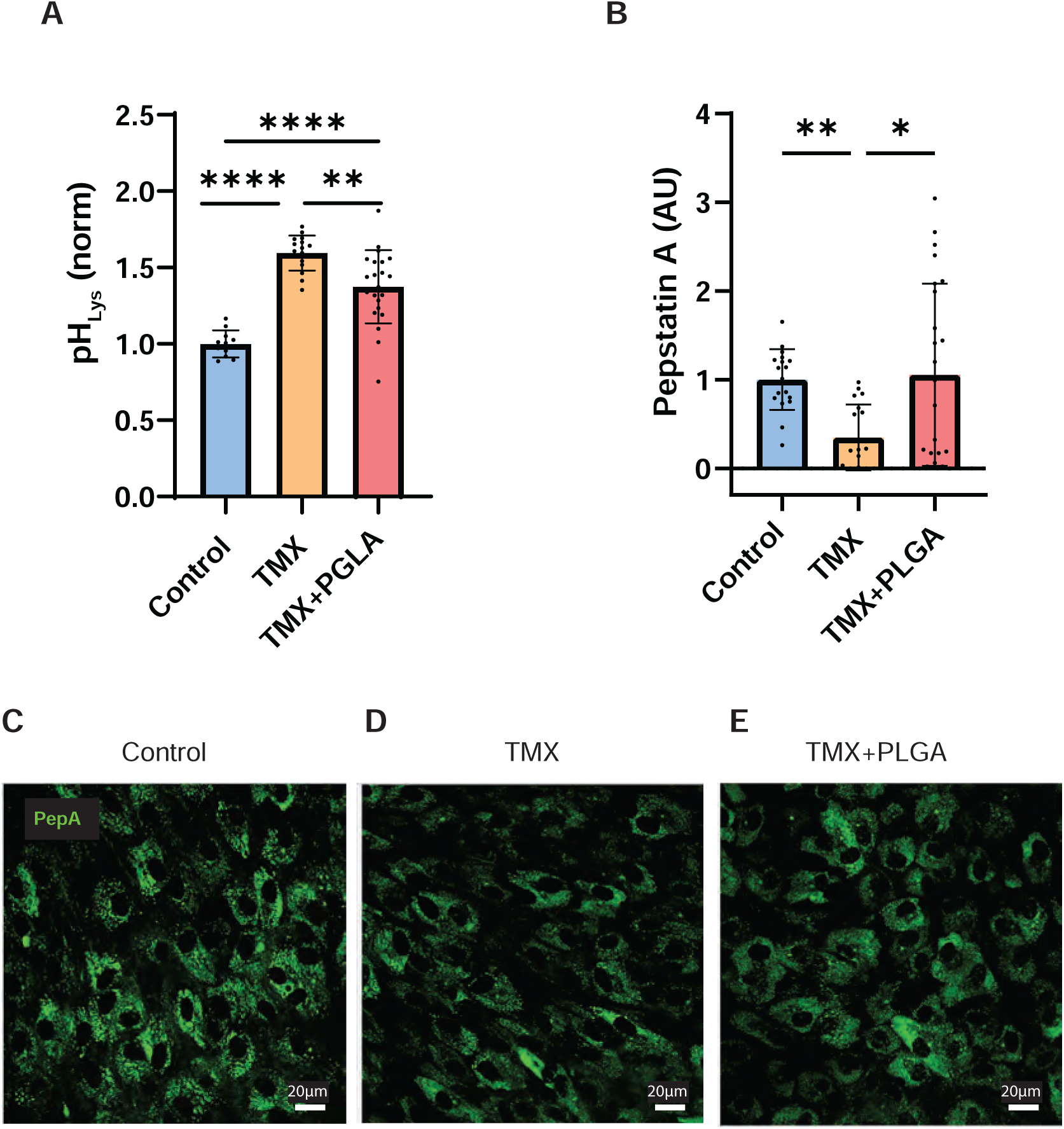
PLGA nanoparticles preserve lysosomal acidity and cathepsin D activity in hONH astrocytes. A. Tamoxifen elevated lysosomal pH in astrocytes, which was prevented by PLGA nanoparticles. **p = 0.0013, ****p < 0.0001 (One-way ANOVA with Tukey’s test; n = 11–22 from 2 trials). Values normalized to daily controls based on ratiometric imaging with LysoSensor Yellow/Blue (340/380 nm excitation; >527 nm emission). A. Bodipy-FL Pepstatin A signal, an indicator of cathepsin D activity, was reduced by tamoxifen and restored with PLGA nanoparticle pre-treatment. *p=0.0177, **p=0.0017 (Kruskal-Wallis with Dunn’s test; n = 18–24 from 3 trials). AU= Arbitrary Units. C–E. Representative images showing Bodipy-FL Pepstatin A fluorescence under control conditions (C), tamoxifen exposure (D), and PLGA nanoparticle + tamoxifen treatment (E).

To assess lysosomal function, the fluorescent probe BODIPY FL-Pepstatin A was used. This probe binds to the active catalytic site of the lysosomal protease Cathepsin D, providing a spatially restricted indicator of lysosomal activity, as both enzyme maturation and probe access are pH-dependent. Tamoxifen treatment reduced BODIPY FL-Pepstatin A fluorescence, indicating impaired lysosomal function. However, this reduction was prevented by PLGA nanoparticle pretreatment (Fig. 7B–E). Strong colocalization between the BODIPY FL-Pepstatin A and LysoTracker Red signals confirmed the lysosomal origin of the fluorescence (Fig. S2).

## Discussion

This study characterized the parameters regulating the delivery of PLGA-based nanoparticles to lysosomes and demonstrated the ability of these nanoparticles to reduce elevated lysosomal pH in epithelial cells and astrocytes. Nanoparticle synthesis from PLGA polymers covalently bound to Cy3 provided a stable fluorescent label to monitor long-term trafficking to lysosomes and optimization of delivery conditions. Quantitative analysis indicated >90% of these nanoparticles localized to lysosomes, suggesting that their chosen size and composition promoted targeted delivery. Enhanced delivery to iPS-RPE cells indicated an advantage over previous data obtained with the ARPE-19 cell line [17], while findings in astrocytes identifies new potential targets.

The sustained presence of these Cy3-labeled nanoparticles in iPS-RPE cells for up to 28 days, without any significant change in size or quantity, supports their potential for extended studies. The ability of the nanoparticles to reacidify lysosomal pH and enhance levels of degradative cathepsin enzymes further supports their functional integration following uptake and lysosomal trafficking. Overall, Cy3-PLGA nanoparticles enabled robust lysosomal targeting and prolonged intracellular residence, underscoring their promise for future *in vivo* applications.

More than 90% of Cy3-PLGA nanoparticles colocalized with lysosomes, suggesting a high specificity for lysosomal targeting. These nanoparticles ranged from 230-280 nm in diameter; while dogma suggests particles with a range of sizes will be delivered to the lysosome, this specific size appears particularly effective [47, 48]. Lysosomal localization was defined as an overlap between green and red fluorescent signals in cells treated with LysoTracker Green. Although acidification by the nanoparticles could theoretically have caused other vesicles to fluoresce green, most green vesicles did not overlap with red fluorescence, and not all Cy3-positive nanoparticles overlapped with green organelles, supporting the independence of the characterization. Functional effects on lysosomal activity also reinforce this localization. Additional characterizations using proteins associated with lysosomes would provide a more definitive confirmation. A recent study using DNA paint found LAMP2 colocalized with 95% of Lamp1-positive vesicles in ARPE19 cells, suggesting either protein would be a reliable marker for future confirmation [49].

Uptake of nanoparticles varied considerably between cells, with some displaying intense fluorescence while others displayed little or none. This variability is particularly notable as the cells used in a given experimental set were derived from the same iPS donor cells. Heterogeneity among neighboring RPE cells has been reported [50–52], though the underlying causes remain unclear. These may reflect a division of labor among cells in a syncytium or be driven by environmental or epigenetic factors. The application of spatial genomics should shed some light on this variability and explain differences in nanoparticle uptake.

While the imaging and analysis methods used here effectively detected Cy3-labeled nanoparticles, they are constrained by the limits of fluorescence microscopy [53]. The size of the nanoparticles approaches this limit, and therefore, size estimates should be interpreted with caution. However, the mean particle size measured over 1-28 days post-loading (∼305nm) was remarkably close to the data obtained with the Zetasizer. This adds rigor to the microscopic detection, and suggests most fluorescent signals represent individual nanoparticles rather than aggregates.

The basic concept of using acidic nanoparticles to restore lysosomal acidity to RPE cells was initially proposed by Baltazar *et al.* [17]. This study substantially advances that work. First, the covalent bonding of Cy3 to PLGA polymers enabled more stable labeling and long-term tracking, improving size and retention measurements. While labeling with Nile-Red enabled the initial characterization, subsequent trials found the dye diffused, complicating localization. Second, the use of iPS-RPE cells offers physiological advances over ARPE19 cells with direct implications for the delivery of nanoparticles to the lysosomes of RPE cells, including enhanced phagocytosis and degradation [34]. While previous work found PLA-based nanoparticles more effective in ARPE-19 cells, this study indicates that PLGA -based nanoparticles are preferable in iPS-RPE cells. The shift may be due to the smaller nanoparticle size or subtle synthesis differences.

This is the first evidence that PLGA-nanoparticles can target lysosomes in optic nerve head astrocytes and restore both an acidic pH and cathepsin D activity in compromised lysosomes. Elevated lysosomal pH has been implicated in astrocyte activation and neuroinflammation [54, 55]. In the optic nerve head, impaired mitophagy was recently associated with retinal degeneration [56]. Larger, more concentrated lipofuscin deposits have been observed in neuroglial cells within axon bundles of glaucomatous patients [57]. Astrocytes carrying glaucoma-associated OPTN mutations showed disrupted autophagy and reduced neurite length [58], while in some models, autophagy inhibition is protective [59]. Acid nanoparticles provide a powerful tool to investigate the role of lysosomal acidification in neurodegeneration.

Acidic nanoparticles are increasingly recognized for their ability to lower lysosomal pH [60]. While pharmacological approaches have advantages in a translational setting, acidic nanoparticles provide targeted, mechanism-specific action, strengthening the case for elevated lysosomal pH as a pathological contributor in preclinical models. Previous studies showed that acidic nanoparticles restored normal lysosomal proteolysis and Ca^2+^ homeostasis in presenilin 1 KO cells [61], and enhanced degradation of phagocytosed debris in cortical astrocytes [62]. Acidic nanoparticles were shown to reduce α-synuclein pathology in mice with human Lewy body extracts [63]. Future *in vivo* trials will indicate the protective potential of acidic nanoparticles in RPE cells and optic nerve head astrocytes under conditions of lysosomal dysfunction.

## Supplemental Figures

**Figure S1.** Nile Red-labeled PLGA nanoparticles (1 mg/ml) incubated with hONH astrocytes overnight. LysoTracker Green stained lysosomes, and Hoechst-stained nuclei. Dye diffusion observed throughout cytoplasm (arrow).

**Figure S2.** Confocal image illustrating variability in nanoparticle uptake in iPS-RPE cells, with neighboring cells displaying different uptake levels.

**Figure S3.** Colocalization of Bodipy FL-Pepstatin A (green) and LysoTracker Red confirming predominantly lysosomal localization. Nuclear stained with Hoechst (blue).

## References

1. Peng W, Minakaki G, Nguyen M, Krainc D. Preserving lysosomal function in the aging brain: insights from neurodegeneration. Neurotherapeutics.16:611–34, 2019.

2. Nixon RA. The aging lysosome: An essential catalyst for late-onset neurodegenerative diseases. Biochim Biophys Acta Proteins Proteom.1868:140443, 2020.

3. Barrett A. Protinases in mammalian cells and tissues. Dingle JT, editor. New York: Elsiver/North-Hollard Biomedical press; 1977. p 220-224.

4. Ishida Y, Nayak S, Mindell JA, Grabe M. A model of lysosomal pH regulation. J Gen Physiol.141:705–20, 2013.

5. Van Dyke RW. Acidification of rat liver lysosomes: quantitation and comparison with endosomes. Am J Physiol-Cell Physiol. 265:C901–C17, 1993.

6. Song W, Wang F, Savini M, Ake A, di Ronza A, Sardiello M, Segatori L. TFEB regulates lysosomal proteostasis. Human Mol Genetics.22:1994-2009, 2013.

7. Yuan S, Jiang SC, Zhang ZW, Fu YF, Yang XY, Li ZL, Hu J. Rethinking of Alzheimer’s disease: Lysosomal overloading and dietary therapy. Front Aging Neurosci.15:1130658, 2023.

8. Wallings RL, Humble SW, Ward ME, Wade-Martins R. Lysosomal dysfunction at the centre of Parkinson’s Disease and Frontotemporal Dementia/Amyotrophic Lateral Sclerosis. Trends Neurosci.42:899–912, 2019.

9. Boya P, Kaarniranta K, Handa JT, Sinha D. Lysosomes in retinal health and disease. Trends Neurosci.46:1067–82, 2023.

10. Guha S, Baltazar GC, Tu LA, Liu J, Lim JC, Lu W, Argall A, Boesze-Battaglia K, Laties AM, Mitchell CH. Stimulation of the D5 dopamine receptor acidifies the lysosomal pH of retinal pigmented epithelial cells and decreases accumulation of autofluorescent photoreceptor debris. J Neurochem.122:823–33, 2012.

11. Lu W, Campagno KE, Tso HY, Cenaj A, Laties AM, Carlsson LG, Mitchell CH. Oral delivery of the P2Y12 receptor antagonist ticagrelor prevents loss of photoreceptors in an ABCA4-/-mouse model of retinal degeneration. Invest Ophthalmol Vis Sci.60:3046–53, 2019.

12. Zeng J, Shirihai OS, Grinstaff MW. Modulating lysosomal pH: a molecular and nanoscale materials design perspective. J Life Sci (Westlake Village).2:25–37, 2020.

13. Pastor-Soler NM, Hallows KR, Smolak C, Gong F, Brown D, Breton S. Alkaline pH- and cAMP-induced V-ATPase membrane accumulation is mediated by protein kinase A in epididymal clear cells. Am J Physiol-Cell Physiol.294:C488–C94, 2008.

14. Guo X, Zuo X, Zhou Z, Gu Y, Zheng H, Wang X, Wang G, Xu C, Wang F. PLGA-Based micro/nanoparticles: an overview of their applications in respiratory diseases. Int J Mol Sci.242023.

15. Panyam J, Labhasetwar V. Biodegradable nanoparticles for drug and gene delivery to cells and tissue. Advanced Drug Delivery Rev.55:329–47, 2003.

16. Félix Lanao RP, Jonker AM, Wolke JG, Jansen JA, van Hest JC, Leeuwenburgh SC. Physicochemical properties and applications of poly(lactic-co-glycolic acid) for use in bone regeneration. Tissue Eng Part B Rev.19:380–90, 2013.

17. Baltazar GC, Guha S, Boesze-Battaglia K, Laties AM, Tyagi P, Kompella UB, Mitchell CH. Acidic nanoparticles restore lysosomal pH and degradative function in compromised RPE cells. PloS One.7:e49635,PMID: 23272048 PMC3525582 2012.

18. Yoo JW, Mitragotri S. Polymer particles that switch shape in response to a stimulus. Proc Natl Acad Sci U S A.107:11205–10, 2010.

19. Panyam J, Zhou WZ, Prabha S, Sahoo SK, Labhasetwar V. Rapid endo-lysosomal escape of poly(DL-lactide-co-glycolide) nanoparticles: implications for drug and gene delivery. Faseb J.16:1217–26, 2002.

20. Kompella UB, Amrite AC, Pacha Ravi R, Durazo SA. Nanomedicines for back of the eye drug delivery, gene delivery, and imaging. Prog Retin Eye Res.36:172–98, 2013.

21. Vargas JA, Finnemann SC. Probing photoreceptor outer segment phagocytosis by the RPE in vivo: models and methodologies. Inter J Mol Sci. 23(7), 2022.

22. Sinha D, Valapala M, Shang P, Hose S, Grebe R, Lutty GA, Zigler JS, Jr., Kaarniranta K, Handa JT. Lysosomes: Regulators of autophagy in the retinal pigmented epithelium. Exp Eye Res.144:46–53, 2016.

23. Valapala M, Wilson C, Hose S, Bhutto IA, Grebe R, Dong A, Greenbaum S, Gu L, Sengupta S, Cano M, Hackett S, Xu G, Lutty GA, Dong L, Sergeev Y, Handa JT, Campochiaro P, Wawrousek E, Zigler JS, Jr., Sinha D. Lysosomal-mediated waste clearance in retinal pigment epithelial cells is regulated by CRYBA1/βA3/A1-crystallin via V-ATPase-MTORC1 signaling. Autophagy.10:480–96, 2014.

24. Kaarniranta K, Blasiak J, Liton P, Boulton M, Klionsky DJ, Sinha D. Autophagy in age-related macular degeneration. Autophagy.19:388–400, 2023.

25. Huang H, Zeng J, Yu X, Du H, Wen C, Mao Y, Tang H, Kuang X, Liu W, Yu H, Liu H, Li B, Long C, Yan J, Shen H. Establishing chronic models of age-related macular degeneration via long-term iron ion overload. American Journal of Physiology-Cell Physiology.326:C1367–C83, 2024.

26. Zhang KR, Jankowski CSR, Marshall R, Nair R, Más Gómez N, Alnemri A, Liu Y, Erler E, Ferrante J, Song Y, Bell BA, Baumann BH, Sterling J, Anderson B, Foshe S, Roof J, Fazelinia H, Spruce LA, Chuang JZ, Sung CH, Dhingra A, Boesze-Battaglia K, Chavali VRM, Rabinowitz JD, Mitchell CH, Dunaief JL. Oxidative stress induces lysosomal membrane permeabilization and ceramide accumulation in retinal pigment epithelial cells. Dis Model Mech.162023.

27. Karema-Jokinen V, Koskela A, Hytti M, Hongisto H, Viheriälä T, Liukkonen M, Torsti T, Skottman H, Kauppinen A, Nymark S, Kaarniranta K. Crosstalk of protein clearance, inflammasome, and Ca(2+) channels in retinal pigment epithelium derived from age-related macular degeneration patients. J Biol Chem.299:104770, 2023.

28. Lakkaraju A, Finnemann SC, Rodriguez-Boulan E. The lipofuscin fluorophore A2E perturbs cholesterol metabolism in retinal pigment epithelial cells. Proc Natl Acad Sci U S A.104:11026–31, 2007.

29. Bergmann M, Schütt F, Holz FG, Kopitz J. Inhibition of the ATP-driven proton pump in RPE lysosomes by the major lipofuscin fluorophore A2-E may contribute to the pathogenesis of age-related macular degeneration. Faseb J.18:562–4, 2004.

30. Liu J, Lu W, Reigada D, Nguyen J, Laties AM, Mitchell CH. Restoration of lysosomal pH in RPE cells from cultured human and ABCA4(-/-) mice: pharmacologic approaches and functional recovery. Invest Ophthalmol Vis Sci.49:772–80, 2008.

31. Choi JA, Seo BR, Koh JY, Yoon YH. Protective effect of zinc against A2E-induced toxicity in ARPE-19 cells: Possible involvement of lysosomal acidification. Heliyon.10:e39100, 2024.

32. Korkka I, Viheriälä T, Juuti-Uusitalo K, Uusitalo-Järvinen H, Skottman H, Hyttinen J, Nymark S. Functional voltage-gated calcium channels are present in human embryonic stem cell-derived retinal pigment epithelium. Stem Cells Transl Med.8:179–93, 2019.

33. Kokkinaki M, Sahibzada N, Golestaneh N. Human induced pluripotent stem-derived retinal pigment epithelium (RPE) cells exhibit ion transport, membrane potential, polarized vascular endothelial growth factor secretion, and gene expression pattern similar to native RPE. Stem Cells.29:825–35, 2011.

34. Fisher CR, Ebeling MC, Geng Z, Kapphahn RJ, Roehrich H, Montezuma SR, Dutton JR, Ferrington DA. Human iPSC- and primary-retinal pigment epithelial cells for modeling age-related macular degeneration. Antioxidants (Basel).112022.

35. Zhang KR, Jankowski CSR, Marshall R, Nair R, Más Gómez N, Alnemri A, Liu Y, Erler E, Ferrante J, Song Y, Bell BA, Baumann BH, Sterling J, Anderson B, Foshe S, Roof J, Fazelinia H, Spruce LA, Chuang JZ, Sung CH, Dhingra A, Boesze-Battaglia K, Chavali VRM, Rabinowitz JD, Mitchell CH, Dunaief JL. Oxidative stress induces lysosomal membrane permeabilization and ceramide accumulation in retinal pigment epithelial cells. Dis Model Mech.162023.

36. Lopez NN, Clark AF, Tovar-Vidales T. Isolation and characterization of human optic nerve head astrocytes and lamina cribrosa cells. Exp Eye Res.197:108103, 2020.

37. Schneider CA, Rasband WS, Eliceiri KW. NIH Image to ImageJ: 25 years of image analysis. Nat Methods.9:671–5, 2012.

38. Liu J, Lu W, Guha S, Baltazar GC, Coffey EE, Laties AM, R.C. R, Reenstra WW, Mitchell CH. Cystic fibrosis transmembrane conductance regulator (CFTR) contributes to reacidification of alkalinized lysosomes in RPE cells Am J Physiol Cell Physiol.303:C160–C169, 2012.

39. Guha S, Coffey EE, Lu W, Lim JC, Beckel JM, Laties AM, Boesze-Battaglia K, Mitchell CH. Approaches for detecting lysosomal alkalinization and impaired degradation in fresh and cultured RPE cells: evidence for a role in retinal degenerations. Exp Eye Res.126:68–76, 2014.

40. Foroozandeh P, Aziz AA. Insight into cellular uptake and intracellular trafficking of nanoparticles. Nanoscale Res Lett.13:339, 2018.

41. Shao X, Guha S, Lu W, Campagno KE, Beckel JM, Mills JA, Yang W, Mitchell CH. Polarized cytokine release triggered by P2X7 receptor from retinal pigmented epithelial cells dependent on calcium iInflux. Cells.9:2537, 2020.

42. Kwon W, Freeman SA. Phagocytosis by the retinal pigment epithelium: recognition, resolution, recycling. Front Immunol. 11:604205, 2020.

43. Lu W, Albalawi F, Beckel JM, Lim JC, Laties AM, Mitchell CH. The P2X7 receptor links mechanical strain to cytokine IL-6 up-regulation and release in neurons and astrocytes. J Neurochem.141:436–48, 2017.

44. Colacurcio DJ, Nixon RA. Disorders of lysosomal acidification-The emerging role of v-ATPase in aging and neurodegenerative disease. Ageing Res Rev.32:75–88, 2016.

45. Altan N, Chen Y, Schindler M, Simon SM. Tamoxifen inhibits acidification in cells independent of the estrogen receptor. Proc Natl Acad Sci U S A.96:4432–7, 1999.

46. Chen Y, Schindler M, Simon SM. A mechanism for tamoxifen-mediated inhibition of acidification. J Biol Chem.274:18364–73, 1999.

47. Kaja S, Payne AJ, Patel KR, Naumchuk Y, Koulen P. Differential subcellular Ca2+ signaling in a highly specialized subpopulation of astrocytes. Exp Neurol.265:59–68, 2015.

48. Baranov MV, Kumar M, Sacanna S, Thutupalli S, van den Bogaart G. Modulation of Immune responses by particle size and shape. Front Immunol.11:607945, 2020.

49. Moreno-Mendieta S, Guillén D, Vasquez-Martínez N, Hernández-Pando R, Sánchez S, Rodríguez-Sanoja R. Understanding the phagocytosis of particles: the key for rational design of vaccines and therapeutics. Pharmaceut Res.39:1823–49, 2022.

50. Bond C, Hugelier S, Xing J, Sorokina EM, Lakadamyali M. Heterogeneity of late endosome/lysosomes shown by multiplexed DNA-PAINT imaging. J Cell Biol.224: e202403116, 2025.

51. Burke JM, Hjelmeland LM. Mosaicism of the retinal pigment epithelium: seeing the small picture. Mol Interv.5:241–9, 2005.

52. Farjood F, Manos JD, Wang Y, Williams AL, Zhao C, Borden S, Alam N, Prusky G, Temple S, Stern JH, Boles NC. Identifying biomarkers of heterogeneity and transplantation efficacy in retinal pigment epithelial cells. J Exp Med.220: e20230913, 2023.

53. Ortolan D, Sharma R, Volkov A, Maminishkis A, Hotaling NA, Huryn LA, Cukras C, Di Marco S, Bisti S, Bharti K. Single-cell–resolution map of human retinal pigment epithelium helps discover subpopulations with differential disease sensitivity. Proc Nat Acad Sci.119:e2117553119, 2022.

54. MacDonald L, Baldini G, Storrie B. Does super-resolution fluorescence microscopy obsolete previous microscopic approaches to protein co-localization? Methods Mol Biol.1270:255–75, 2015.

55. Zeng J, Indajang J, Pitt D, Lo CH. Lysosomal acidification impairment in astrocyte-mediated neuroinflammation. J Neuroinflam.22:72, 2025.

56. Lo CH, Zeng J. Defective lysosomal acidification: a new prognostic marker and therapeutic target for neurodegenerative diseases. Trans Neurodegen.12:29, 2023.

57. Yazdankhah M, Ghosh S, Liu H, Hose S, Zigler JS, Jr., Sinha D. Mitophagy in astrocytes is required for the health of optic nerve. Cells.12; 2496, 2023.

58. Fernandez de Castro JP, Mullins RF, Manea AM, Hernandez J, Wallen T, Kuehn MH. Lipofuscin in human glaucomatous optic nerves. Exp Eye Res.111:61–6, 2013.

59. Gomes C, VanderWall KB, Pan Y, Lu X, Lavekar SS, Huang K-C, Fligor CM, Harkin J, Zhang C, Cummins TR, Meyer JS. Astrocytes modulate neurodegenerative phenotypes associated with glaucoma in OPTN(E50K) human stem cell-derived retinal ganglion cells. Stem Cell Reports.17:1636–49, 2022.

60. Dixon A, Shim MS, Nettesheim A, Coyne A, Su C-C, Gong H, Liton PB. Autophagy deficiency protects against ocular hypertension and neurodegeneration in experimental and spontaneous glaucoma mouse models. Cell Death & Disease.14:554, 2023.

61. Bourdenx M, Daniel J, Genin E, Soria FN, Blanchard-Desce M, Bezard E, Dehay B. Nanoparticles restore lysosomal acidification defects: Implications for Parkinson and other lysosomal-related diseases. Autophagy.12:472–83, 2016.

62. 58. Lee JH, McBrayer MK, Wolfe DM, Haslett LJ, Kumar A, Sato Y, Lie PP, Mohan P, Coffey EE, Kompella U, Mitchell CH, Lloyd-Evans E, Nixon RA. Presenilin 1 maintains lysosomal Ca2+ homeostasis via TRPML1 by regulating vATPase-mediated lysosome acidification. Cell Rep.12:1430–44, 2015.

63. Lööv C, Mitchell CH, Simonsson M, Erlandsson A. Slow degradation in phagocytic astrocytes can be enhanced by lysosomal acidification. Glia.63:1997–2009, 2015.

64. Arotcarena ML, Soria FN, Cunha A, Doudnikoff E, Prévot G, Daniel J, Blanchard-Desce M, Barthélémy P, Bezard E, Crauste-Manciet S, Dehay B. Acidic nanoparticles protect against α-synuclein-induced neurodegeneration through the restoration of lysosomal function. Aging Cell.21:e13584, 2022.

