## Supplemental data and files for "Sustained Lysosomal Delivery of Enhanced Cy3-Labeled Acid Nanoparticles Restores Lysosomal pH in Retinal Pigment Epithelial Cells and Astrocytes"

### Supplemental Figures

**Figure S1**. Nile Red-labeled PLGA nanoparticles (1 mg/ml) incubated with hONH astrocytes overnight. LysoTracker Green stained lysosomes, and Hoechst-stained nuclei. Dye diffusion observed throughout cytoplasm (arrow).

**Figure S2.** Confocal image illustrating variability in nanoparticle uptake in iPS-RPE cells, with neighboring cells displaying different uptake levels.

**Figure S3.** Colocalization of Bodipy FL-Pepstatin A (green) and LysoTracker Red confirming predominantly lysosomal localization. Nuclear stained with Hoechst (blue).

Figure S1

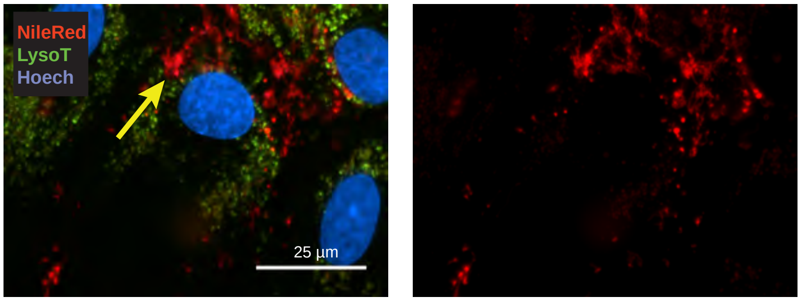

Figure S2

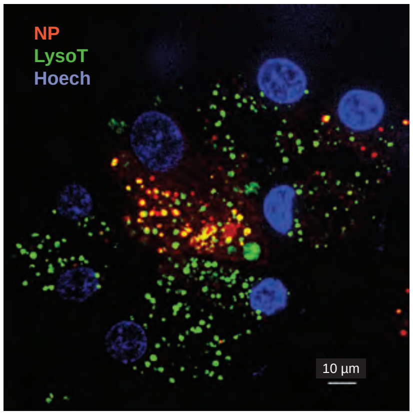

Figure S3

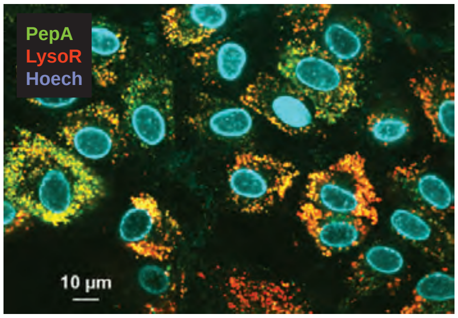
